# Loss of Foxc1 and Foxc2 function in chondroprogenitor cells disrupts endochondral ossification

**DOI:** 10.1101/2021.01.27.428508

**Authors:** Asra Almubarak, Rotem Lavy, Nikola Srnic, Yawen Hu, Devi P. Maripuri, Tsutomo Kume, Fred B Berry

## Abstract

Endochondral ossification forms and grows the majority of the mammalian skeleton and is tightly controlled through gene regulatory networks. The forkhead box transcription factors *Foxc1* and *Foxc2* have been demonstrated to regulate aspects of osteoblast function in the formation of the skeleton but their roles in chondrocytes to control endochondral ossification are less clear. We demonstrate that Foxc1 expression is directly regulated by SOX9 activity, one of the earliest transcription factors to specify the chondrocyte lineages. Moreover we demonstrate that elevelated expression of Foxc1 promotes chondrocyte differentiation in mouse embryonic stem cells and loss of Foxc1 function inhibits chondrogenesis in vitro. Using chondrocyte-targeted deletion of Foxc1 and Foxc2 in mice, we reveal a role for these factors in chondrocyte differentiation in vivo. Loss of both Foxc1 and Foxc2 caused a general skeletal dysplasia predominantly affecting the vertebral column. The long bones of the limb were smaller and mineralization was reduced and organization of the growth plate was disrupted. In particular, the stacked columnar organization of the proliferative chondrocyte layer was reduced in size and cell proliferation in growth plate chondrocytes was reduced. Differential gene expression analysis indicated disrupted expression patterns in chondrogenesis and ossification genes throughout the entire process of endochondral ossification in *Col2-cre;Foxc1^Δ/Δ^;Foxc2^Δ/Δ^* embryos. Our results suggest that *Foxc1* and *Foxc2* are required for correct chondrocyte differentiation and function. Loss of both genes results in disorganization of the growth plate, reduced chondrocyte proliferation and delays in chondrocyte hypertrophy that prevents correct ossification of the endochondral skeleton.

## Introduction

The majority of the mammalian skeleton forms through a mechanism known as endochondral ossification (1, 2). In this developmental event, mesenchymal progenitor cells condense at the sites of newly forming bone and differentiate into chondrocytes cells. These chondrocytes undergo continued differentiation and organize themselves into cell layers that form the growth plate which ultimately drives endochondral bone growth. Round resting zone chondrocytes form at the distal ends of long bones. These cells then differentiate inwards to become flattened, highly proliferative columnar chondrocytes. Much of the extension of the bone length is achieved by the proliferative activities of these cells. Columnar chondrocytes exit the cell cycle to differentiate into pre-hypertrophic chondrocytes and then enlarge to form hypertrophic chondrocytes that form a mineralized matrix and sets the foundation for future bone growth. The fate of the hypertrophic chondrocytes is split into a number of outcomes. A portion of these cells will undergo apoptosis and are removed from the bone; alternatively, hypertrophic chondrocytes will transdifferentiate into bone-forming osteoblast cells that contribute to the ossified bone structure (3, 4). Additional osteoblast cells originate from a cell layer, the periosteum, that lines the newly forming bone, and invade into the newly formed marrow spaced along with blood vessels.

The differentiation of chondrocytes to control growth plate functions is a tightly regulated process controlled by multiple signalling networks. In particular, cross talk between Indian Hedgehog (IHH) and Parathyroid Hormone Related Peptide (PTHrP) signals coordinate the proliferation of the columnar chondrocytes and their exit from the cell cycle to differentiate into prehypertrophic and hypertrophic chondrocytes (5, 6). In addition, Fibroblast Growth Factor (FGF) signalling networks also regulate growth plate chondrocyte function needed for proper bone growth (7, 8). Disruption to these pathways can affect the formation of the skeleton and result in bone growth disorders in humans (9).

Campomelic dysplasia (CD) [OMIM # 114290] is a lethal skeleton malformation characterised by bowed limb bones and a reduced size of the rib cage (10). Mutations in the transcription factor gene *SOX9* cause CD and extensive functional analysis of *SOX9* has defined it as a master regulator of the chondrocyte lineage (10–12). SOX9 function regulates multiple stages of chondrocyte differentiation and development including the initial acquisition of the chondrocyte fate, the proliferation of growth plate chondrocytes and the transition to hypertrophic chondrocytes (12–14). In addition to its profound role in directing cells down the chondrocyte lineage, Sox9 acts to prevent differentiation towards other lineages (13, 15, 16). Although important in the formation of the chondrocyte lineage, SOX9 is dispensable for the initiation of the chondrogenic lineage and the induction of gene expression patterns associated with this fate (17). This finding suggests that additional transcription factors function along with SOX9 to control chondrocyte formation during endochondral ossification.

The forkhead box (FOX) transcription factors are candidates for early regulators of chondrocyte differentiation. Both *Foxc1* and *Foxc2* genes are expressed in the condensing mesenchyme of the presumptive endochondral skeleton (18–21). Furthermore, *Foxc1* and *Foxc2* are required for proper endochondral ossification as mice deficient for *Foxc1* or *Foxc2* display disruptions to the formation of the endochondral skeleton (19, 20, 22). Homozygous null *Foxc1* mouse mutants die shortly after birth and display small rib cages that lack an ossified sternum (19). The neural arches of the vertebral column are not fully mineralized in these mutants. The limbs are shorter in *Foxc1* deficient mice and *Foxc1* can regulate IHH signalling to control endochondral growth in the limb (21). *Foxc2* homozygous null mutant mice also die prior to birth with patterning and ossification defects apparent in the axial skeleton (skull, rib cage and vertebral column) (20). Although both *Foxc1* and *Foxc2* are expressed at high levels in chondrocytes of the developing limbs (21), loss of function mutation of either gene results in milder phenotypes than that observed in the axial skeleton. As FOXC1 and FOXC2 proteins have near identical DNA-binding domains (23–25) it is possible that *Foxc1* or *Foxc2* may compensate for the loss of the other. Compound *Foxc1^−/−^;Foxc2^−/−^* mice arrest in development prior to the onset of skeletal formation and therefore the combined functions of these genes in endochondral ossification is not known (26).

We wished to determine how *Foxc1* and *Foxc2* function in chondrocytes to regulate endochondral ossification. We first used in vitro assays to demonstrate that *Foxc1* expression could be directly regulated by SOX9 activity and that gain or loss of *Foxc1* function could positively or negatively regulate in vitro chondrocyte differentiation, respectively. We also demonstrate that the loss of both *Foxc1* and *Foxc2* function in early chondrocyte cells in the developing mouse disrupts normal endochondral ossification processes. These findings indicate that *Foxc1* and *Foxc2* gene function is required in the chondrocyte cells to correctly form the endochondral skeleton.

## Results

### SOX9 directly regulates Foxc1 expression

*Foxc1* and *Foxc2* are expressed in condensing prechondrogenic mesenchyme cells at a time when SOX9 is active (18, 19). Given that their mRNA expression is reduced in Sox9 deficient chondrogenic tissues (17) we sought to determine whether *Foxc1* and *Foxc2* were directly regulated by SOX9. First we used an inducible mouse embryonic stem (mES) cell line that contains a Tet-off incubible *Sox9* gene (27). Upon the removal of doxycycline for 48 hours we observed an elevation of SOX9 protein and mRNA level as well as an increase in *Col2a* mRNA, a known SOX9 target of transcriptional regulation (28). Expression of *Foxc1* was also elevated in response to SOX9 induction but levels of *Foxc2* mRNA remain unchanged (Fig 1A). We next examined SOX9 ChiP-seq data and identified four SOX9 binding peaks near the mouse *Foxc1* gene (29). Three peaks were located upstream of *Foxc1* which we termed Distal A (mm10 chr13:31,764,541-31,764,717), B(mm10 chr13:31,765,465-31,765,623), C (mm10 chr13:31,779,560-31,779,803) and one peak located downstream of *Foxc1* we termed Distal D (mm10 chr13:31,820,626-31,820,791). No peaks were found in proximity to the *Foxc2* gene. We verified SOX9 binding to these sites using Chromatin Immunoprecipitation (ChIP) in ATDC5 cells and found SOX9 was associated with all four regulatory elements as well as the known SOX9 binding site in the intron 1 enhancer of the *Col2a* gene (Fig 1B). Next, we cloned each regulatory region into a luciferase reporter that contains a basal promoter and tested for activation by SOX9 in ATDC5 cells. We found that only Distal C was activated in response to SOX9. Although Distal B was not activated by SOX9 we did detect elevated activity in ATDC5 cells compared to the empty reporter vector suggesting this element may confer Sox9-independent chondrocyte regulatory activity for *Foxc1* expression. Together these findings indicate the *Foxc1* is a direct target of SOX9 transcriptional regulatory activity.

**Figure 1.**
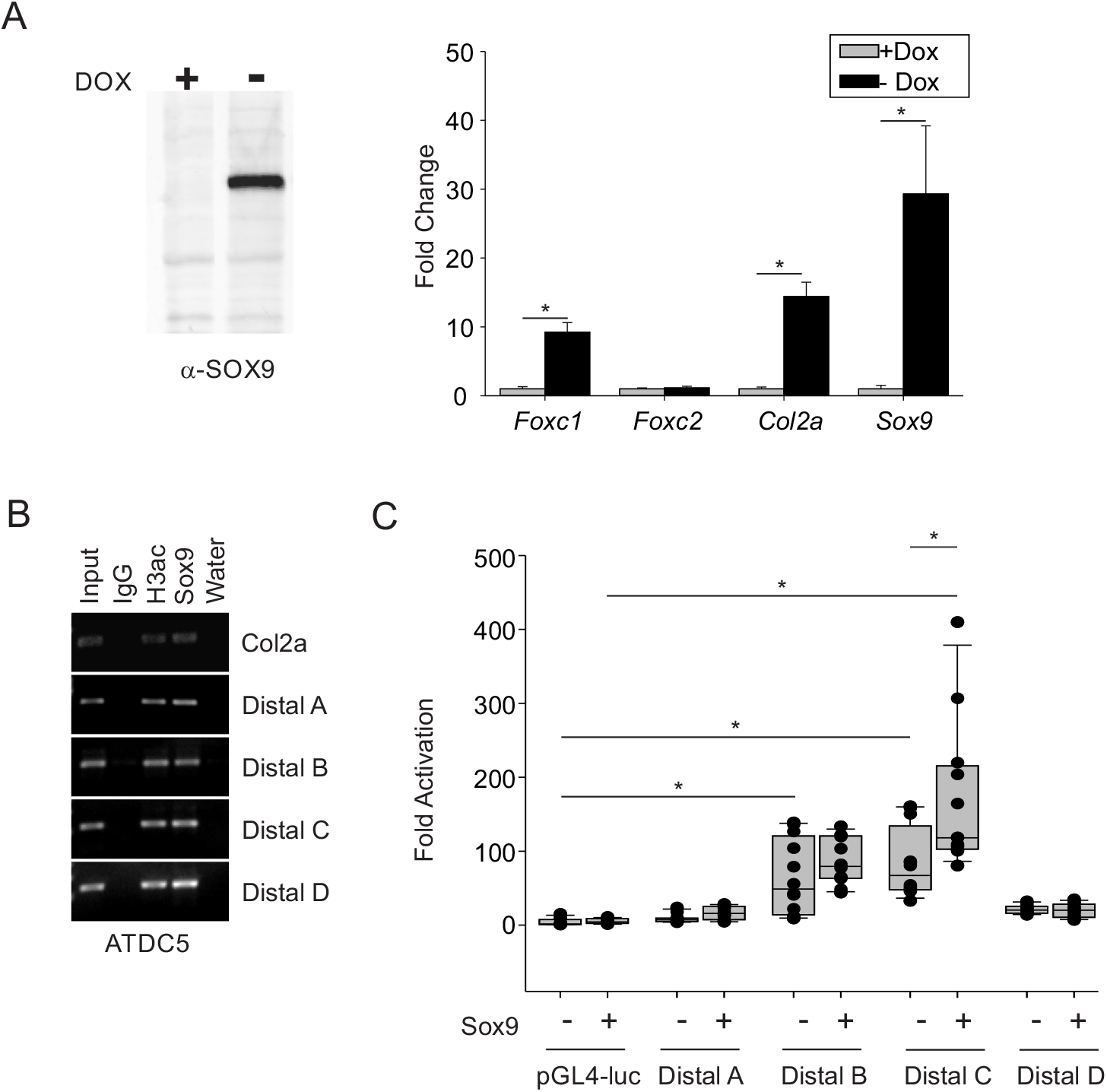
SOX9 regulates expression of *Foxc1*. (A) SOX9 expression was induced in mouse embryonic stem cells containing a doxycycline (Dox)-inducible cassette. Expression was induced for 48h by removal of Dox. Expression of Foxc1, Foxc2 Col2a and Sox9 mRNA was determined by qRT-PCR. Data are presented from three independent experiments. Error bars represent standard deviation. (B) SOX9 binding to regulatory elements in *Col2a* and *Foxc1* was determined by Chromatin Immunoprecipitation (ChIP) in ATDC5 chondrocyte cells. Four Sox9 binding sites in the regulatory region of *Foxc1*(Distal A-D) were identified from previous ChIP-seq experiments (Ohba et al 2015). (C) *Foxc1* Distal regulatory elements were cloned into pGL4-luciferase reporters and transfected along with *Sox9* in ATDC5 cells. Only Foxc1-Distal C was activated by *Sox9*. * p-value <0.05.

### Foxc1 regulates chondrocyte differentiation in vitro

Next we examined whether *Foxc1* functions in regulating chondrocyte differentiation. First we used a *Foxc1*-inducible mES cell line (30) to assess whether *Foxc*1-over expression influences chondrocyte differentiation. We employed a chondrocyte differentiation protocol outlined in Fig 2A and described in (31). Dox was removed from mES cells 2 days prior to chondrocyte differentiation and we confirmed that FOXC1 protein levels were elevated by Dox removal (Fig 2B). Expression o*f Sox9, Sox6, Col2a, Runx2* and *ColX* mRNA was elevated in response to FOXC1 protein induction at 21 days of differentiation compared to uninduced controls. These results indicate that enforced *Foxc1* expression could enhance the differentiation capacity of mES cells.

**Figure 2.**
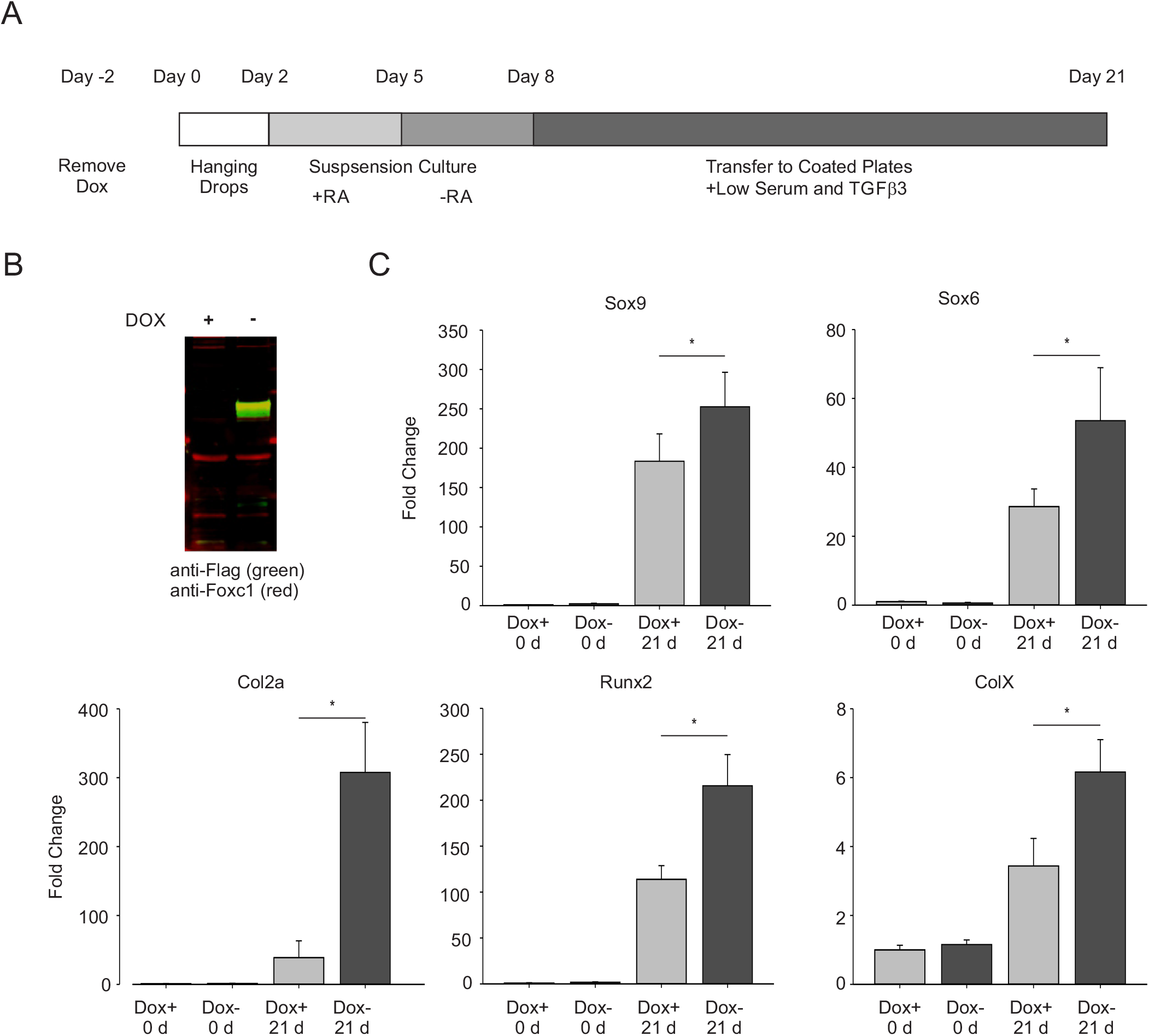
Elevated Foxc1 promotes chondrocyte differentiation of mouse embryonic stem cells (mES). Schematic of chondrocyte differentiation protocol for mES cells (Kawaguchi et al 2005). (B) Expression of FOXC1 was confirmed in mES cells containing a Dox-inducible flag-tagged Foxc1 expression cassette. (C) Chondrocyte differentiation in mES cells was monitored by measuring expression of Sox9, Sox6, Col2a, Runx2 and ColX expression. Data presented are the means from three biological replicates. Error bars represent standard deviation. * p-value <0.05

To examine whether loss of Foxc1 function affected chondrocyte differentiation we used CrisprCas9 to mutate the *Foxc1* gene in ATDC5 cells. We generated a cell line (crFOXC1) that introduced a premature stop codon that would truncate FOXC1 protein after helix 2 in the Forkhead domain. Given that this truncation occurred before helix 3, the DNA recognition helix, this mutation would result in a non-functioning protein. Reduced FOXC1 protein levels were observed in this cell line (Fig 3A). We then induced chondrocyte differentiation by supplementing the culture media with insulin, transferrin and selenium to initiate chondrocyte differentiation. After 21 days of differentiation we detected reduced Alcian blue stained chondrogenic nodules in the crFOXC1 cells (Fig 3B). Furthermore loss of *Foxc1* function reduced levels of genes expressed in early chondrocyte (*Col2a, Sox9*),proliferating chondrocytes (*Fgfr3*), prehypertrophic chondrocytes (*Ihh*) and hypertrophic chondrocytes (*ColX* and *Mmp13*). Together these findings indicate an impairment of chondrocyte differentiation when *Foxc1* function is lost

**Figure 3.**
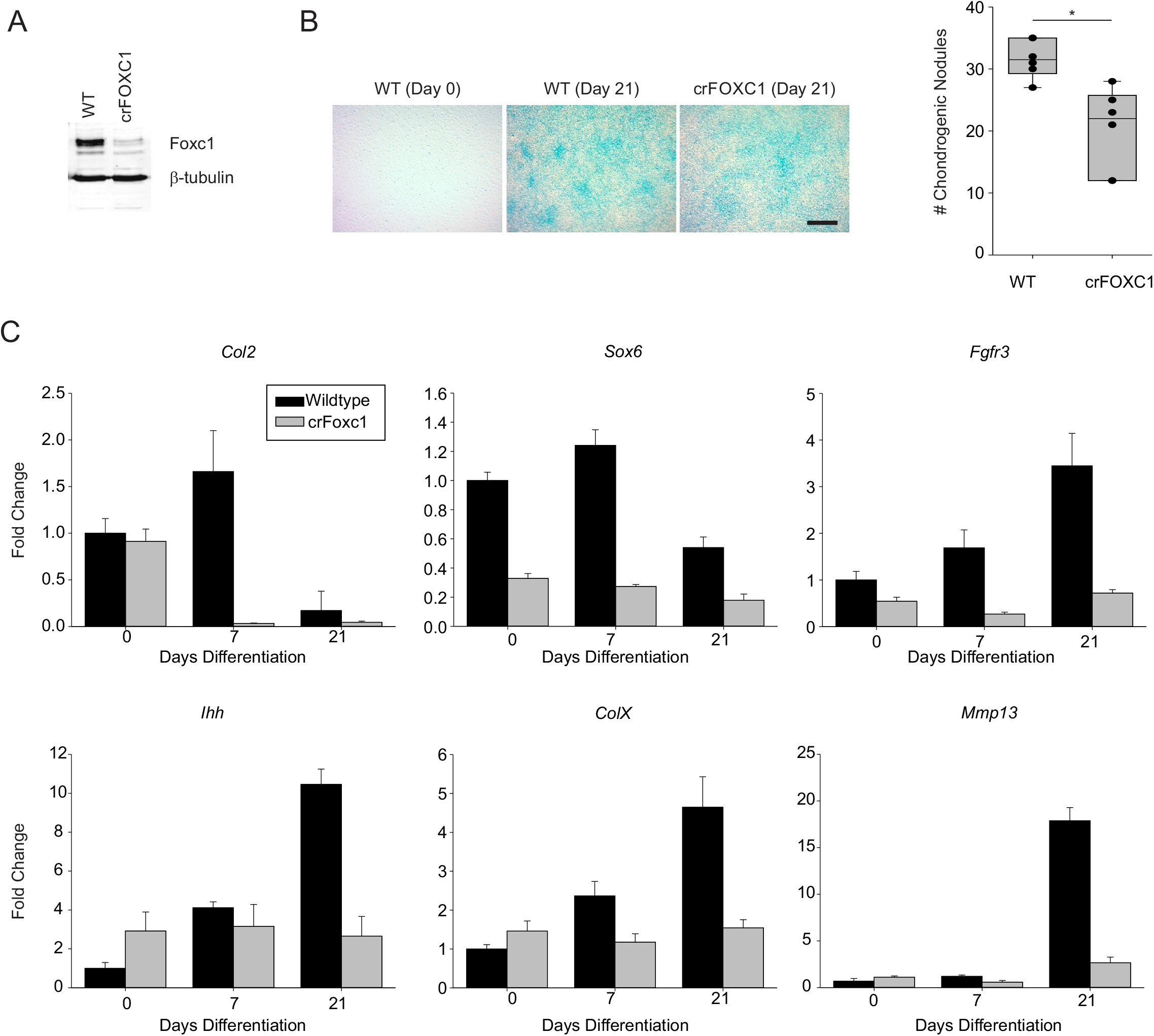
Loss of Foxc1 expression in ATDC5 cells alters chondrocyte differentiation. (A) Deletion of Foxc1 expression was achieved through Crispr mutagenesis. Reduced protein levels observed in Foxc1 mutant cells (crFOXC1). (B) Wild type and crFOXC1 cells were differentiated for 21 days and chondrogenesis measured by Alcian Blue staining. The number of chondrogenic nodules were counted in wild type and mutant cells. (C) Levels of chondrocyte-expressed genes are affected in Foxc1 mutant ATDC5 cells after 21 days of differentiation.

### Foxc1 and Foxc2 are expressed in the perichondrium and the resting zone of the growth plate

Next we examined the localization of *Foxc1* and *Foxc2* expression in the developing skeleton. We focused on both *Foxc1* and *Foxc2* as each gene has a documented function for the proper formation of the skeleton (19–21). More importantly a direct comparison of gene expression patterns of *Foxc1* and *Foxc2* mRNA in the developing skeleton has yet to be performed. We used dual labeling in situ hybridization to simultaneously localize *Foxc1* and *Foxc2* mRNAs in the growth plate of the mouse hindlimb. We observed both overlapping and distinct expression patterns for *Foxc1* and *Foxc2* mRNAs. In the tibia at 14.5 days post coitum (dpc), *Foxc1* mRNA was strongly detected in the perichondrium, resting zone chondrocytes and late hypertrophic chondrocytes/primary ossification center and lower expression detected in the proliferating and prehypertrophic chondrocytes (Fig 4A). We also observed strong *Foxc1* expression in the cells lying between the tibia and femur. *Foxc2* mRNA expression was restricted to the perichondrium and primary ossification center at this time point (Fig 4A). A distinct expression pattern for *Foxc2* mRNA was detected in the outer cell layer of perichondrium. Expression of *Foxc1* and *Foxc2* mRNAs in the proximal tibia at 16.5 dpc continued to be enriched in the surrounding perichondrium (Fig 4B-E). *Foxc1* and *Foxc2* expressing cells were also detected in the resting zone chondrocytes (Fig 4B) and in putative borderline chondrocytes cells found between the proliferating zone chondrocytes and the perichondrium (Fig 4F). Very little *Foxc1* or *Foxc2* mRNA was detected in proliferating chondrocyte or hypertrophic chondrocyte cells (Fig 4G). Together these results indicate that *Foxc1* and *Foxc2* have restricted expression patterns in the developing bones: expression of both *Foxc1* and *Foxc2* is abundant in early stages of endochondral ossification (14.5 dpc) and becomes restricted to the perichondrium and resting chondrocytes at later stages (16.5 dpc).

**Figure 4.**
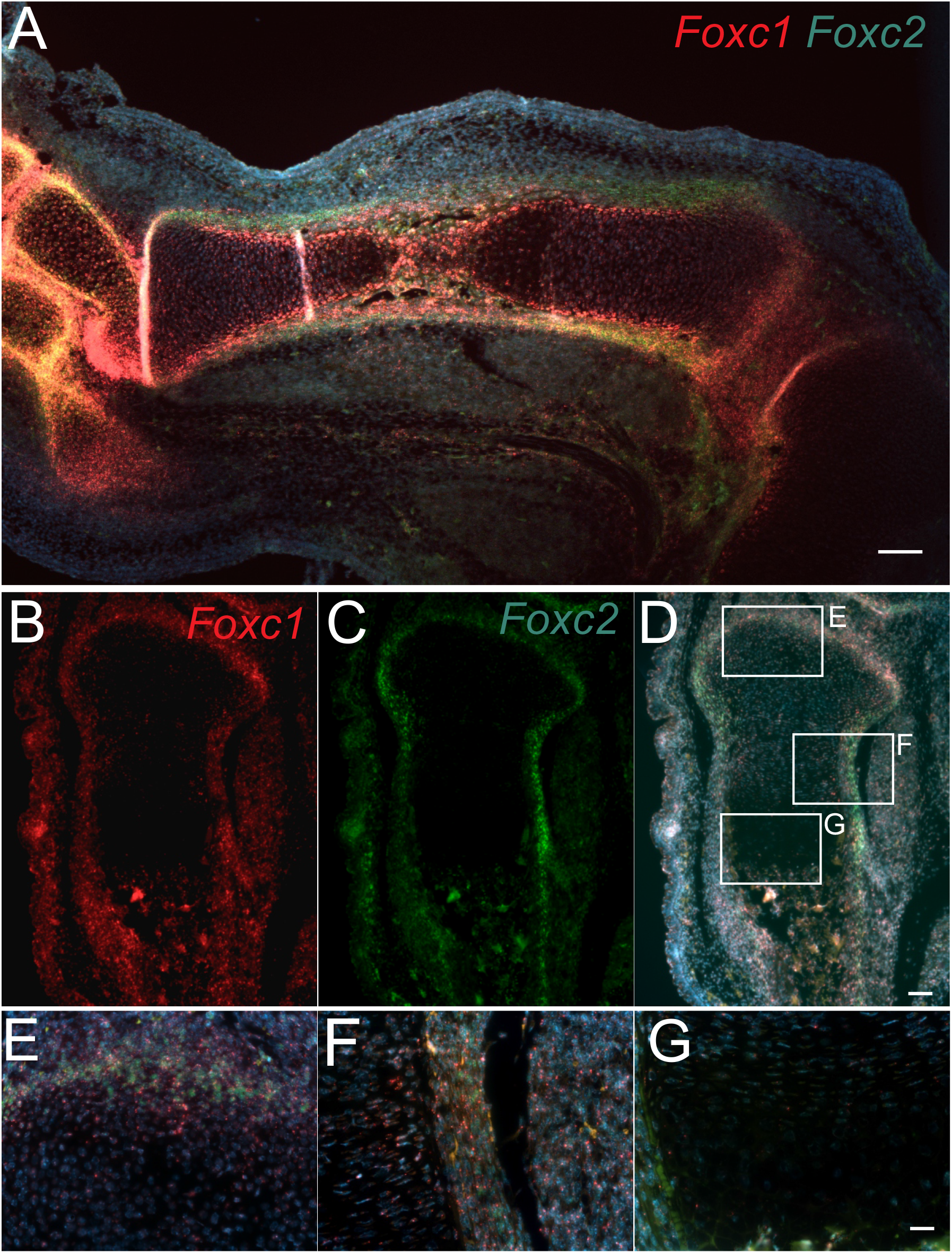
Foxc1 and Foxc2 are expressed in the perichondrium and resting zone of the growth plate. (A) Localization of Foxc1 and Foxc2 expression was determined in the 14.5 dpc hind limb by in situ hybridization. (B)Foxc1 and (C) Foxc2 mRNA expression in the proximal tibia at 16.5 dpc was abundant in the perichondrium. (D). Foxc1 and Foxc2 mRNA displayed both overlapping and distinct expression patterns in the developing limb. (E). Foxc1 transcripts (Red) were detected in the perichondrium and resting zone, while Foxc2(Green) was mainly expressed in the perichondrium. Low levels of Foxc1 and Foxc2 expression were detected in the proliferative zone (F) and hypertrophic zone (G).

### Deletion of Foxc1 and Foxc2 in chondrocyte progenitors causes skeletal dysplasia

In order to understand the roles of *Foxc1* and *Foxc2* in the formation of the endochondral skeleton, we generated conditional, compound *Foxc1* and *Foxc2* mutant mice using the Col2-cre driver strain (32). We chose to delete both *Foxc1* and *Foxc2* genes to eliminate any potential genetic compensation that may occur when one *Foxc* paralog was deleted. First we assessed the cre activity of this strain by crossing *Col2-cre* mice with ROSA26^mTmG^ mice, that expresses EGFP when cre recombinase is present. At 14.5 dpc, EGFP was detected throughout the developing skeleton. In the hindlimb cre activity was detected in the growth plate chondrocytes, the perichondrium and primary ossification center (Fig 5A,B). This expression pattern indicates that the Col-cre strain is active in cells of the developing skeleton and in cell layers that express *Foxc1* and *Foxc2*.

**Figure 5.**
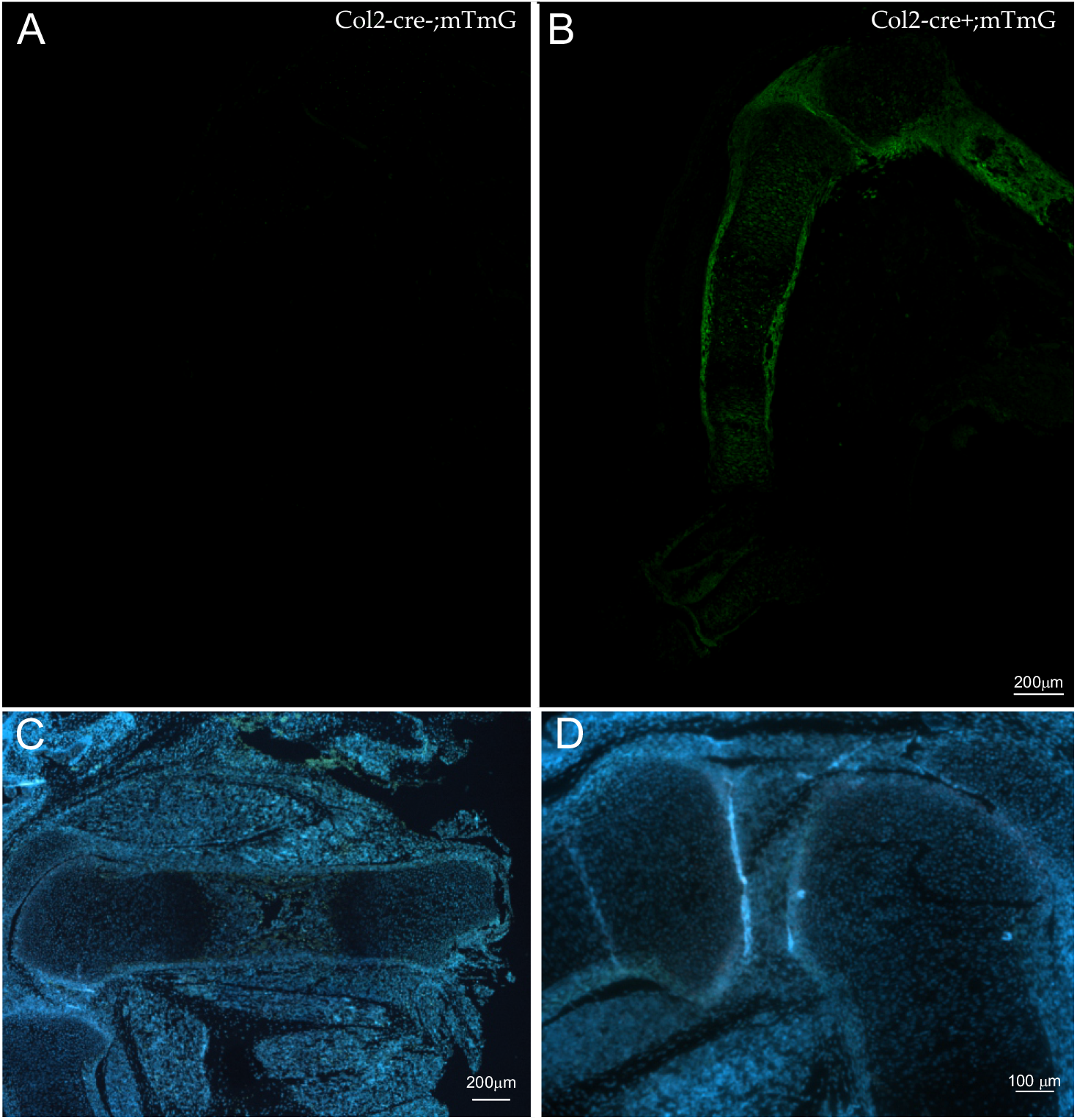
Col2-cre ablation of Foxc1 and Foxc2 expression in the developing limb. (A) (B) ROSA26^tm4(ACTB-tdTomato,-EGFP)^(mTmG) mice were crossed to Col2-Cre mice to monitor Cre activity in the developing limb at 14.5 dpc. EGFP was only detected in limb of mice containing the Col2-cre transgene. (C)(D) To create chondrocyte-specific Foxc1 and Foxc2 mutant mice, Col2-cre mice were crossed with homozygous “floxed” Foxc1 and Foxc2 mice. No expression of Foxc1 and Foxc2 mRNA was detected by in situ hybridization in the developing humerus or proximal tibia in *Col2-cre;Foxc1^Δ/Δ^;Foxc2^Δ/Δ^* mice at 16.5 dpc.

Next we crossed the Col2-cre mice to homozygous floxed (fl) *Foxc1;Foxc2* mice (33) to generate *Col2-cre;Foxc1^+/Δ^;Foxc2^+/Δ^* heterozygotes. These mice were viable and displayed no overt health issues. To delete both copies of *Foxc1* and *Foxc2* in chondrocyte progenitors we crossed male *Col2-cre;Foxc1^+/Δ^;Foxc2^+/Δ^* mice with *Foxc1^fl/fl^;Foxc2^fl/fl^* females. We confirmed by in situ hybridization that *Foxc1* and *Foxc2* mRNA expression was lost in the hindlimbs of 16.5 dpc *Col2-cre;Foxc1^Δ/Δ^;Foxc2^Δ/Δ^* embryos (Fig 5C,D). No viable *Col2-cre;Foxc1^Δ/Δ^;Foxc2^Δ/Δ^, Col2-cre;Foxc1^Δ/Δ^;Foxc2^+/Δ^, Col2-cre;Foxc1^+/Δ^;Foxc2^Δ/Δ^* pups were found at birth. We then isolated embryos at 18.5 dpc for whole skeletal staining and found all genotypes were present at expected Mendelian ratios. We detected a skeletal hypoplasia that worsened when *Foxc* gene dosage was lost (Fig 6A). The skeletons of *Col2-cre;Foxc1^+/Δ^;Foxc2^+/Δ^* embryos at 18.5 dpc formed correctly and displayed no overt phenotypes compared with control (cre negative) embryos. *Col2-cre;Foxc1^+/Δ^;Foxc2^Δ/Δ^* displayed skeletal anomalies including underdeveloped occipital bones and vertebrae. In the cervical vertebrae, the atlas and axis bones were markedly reduced in size compared to control embryos and compound *Col2-cre;Foxc1^+/Δ^;Foxc2^+/Δ^* mice. The skeletons of *Col2-cre;Foxc1^Δ/Δ^;Foxc2^+/Δ^* displayed a smaller, misshapen rib cage, with malformed cervical vertebrae and occipital bones (Fig 6A). The most severe phenotype was observed in the *Col2-cre;Foxc1^Δ/Δ^;Foxc2^Δ/Δ^* embryos and will be the remaining focus of this paper. These embryos displayed a markedly underdeveloped skeleton. The occipital bones were missing, giving the skull a domed appearance (6/6 embryos). The cervical vertebrae were absent in *Col2-cre;Foxc1^Δ/Δ^;Foxc2^Δ/Δ^* embryos and ossification of the remaining bones in the vertebral column was impaired (6/6 embryos; Fig 6B). The rib cage was misshapen but patterned correctly as no missing or fused ribs were detected in the *Col2-cre;Foxc1^Δ/Δ^;Foxc2^Δ/Δ^* embryos (Fig 6B). Likewise, the overall patterning of the limbs appeared normal although ossification was reduced or delayed (Fig 6B,C). Ossification of the proximal and middle phalanges bones was absent in the *Col2-cre;Foxc1^Δ/Δ^;Foxc2^Δ/Δ^* (6/6 embryos). Ossification of the distal phalange bones was variable, with ossification detected in 4 out 6 embryos examined.

**Figure 6.**
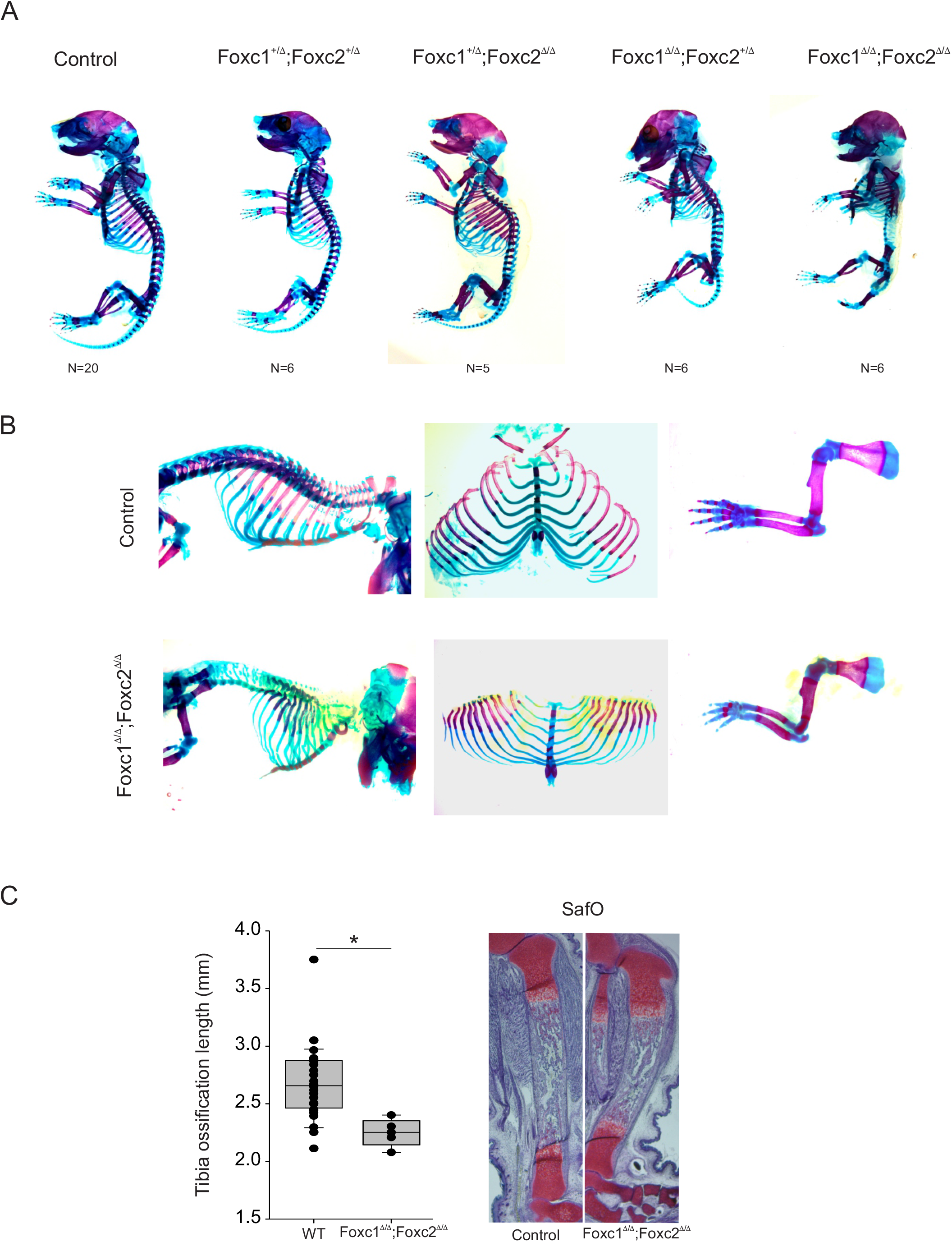
Col2-cre deletion of Foxc1 and Foxc2 causes skeletal hypoplasia. (A). Alizarin red and Alcian blue skeleton preps from 18.5 dpc embryos. Control embryos lack the Col2-cre transgene. (B) Disrupted endochondral ossification can be observed in the axial and appendicular skeleton of *Col2-cre;Foxc1^Δ/Δ^;Foxc2^Δ/Δ^* embryos. (C)Length of ossification is reduced in the tibia at 18.5 dpc of *Col2-cre;Foxc1^Δ/Δ^;Foxc2^Δ/Δ^* mice is reduced.

### Foxc1 and Foxc2 are required for the formation of the growth plate

The limbs of *Col2-cre;Foxc1^Δ/Δ^;Foxc2^Δ/Δ^* embryos displayed reduced mineralization and reduction in length (Fig 6C). We therefore decided to investigate the formation of the growth plate of these mutants to identify the contribution of *Foxc1* and *Foxc2* to the function of this structure. Sections through the 16.5 dpc proximal tibia revealed growth plate anomalies in the *Col2-cre;Foxc1^Δ/Δ^;Foxc2^Δ/Δ^* embryos (Fig 7A). In particular, the characteristic stacked organization of the proliferative zone chondrocytes was not as prevalent in the *Col2-cre;Foxc1^Δ/Δ^;Foxc2^Δ/Δ^* embryos. The length of the resting zone chondrocytes layer was expanded, while the proliferating zone layer was reduced in the *Col2-cre;Foxc1^Δ/Δ^;Foxc2^Δ/Δ^* embryos (Fig 7B). We next assessed whether this reduction in size of the proliferating zone chondrocytes corresponded to a reduction in cell proliferation. As observed in Fig 7C, we did detect a reduction in the number of Ki67-positive cells in the growth plate of *Col2-cre;Foxc1^Δ/Δ^;Foxc2^Δ/Δ^* embryos. We also examined whether cell death was affected in *Col2-cre;Foxc1^Δ/Δ^;Foxc2^Δ/Δ^* embryos and found no changes in the number of apoptotic cells in the growth plates between control and *Col2-cre;Foxc1^Δ/Δ^;Foxc2^Δ/Δ^* embryos (data not shown).

**Figure 7.**
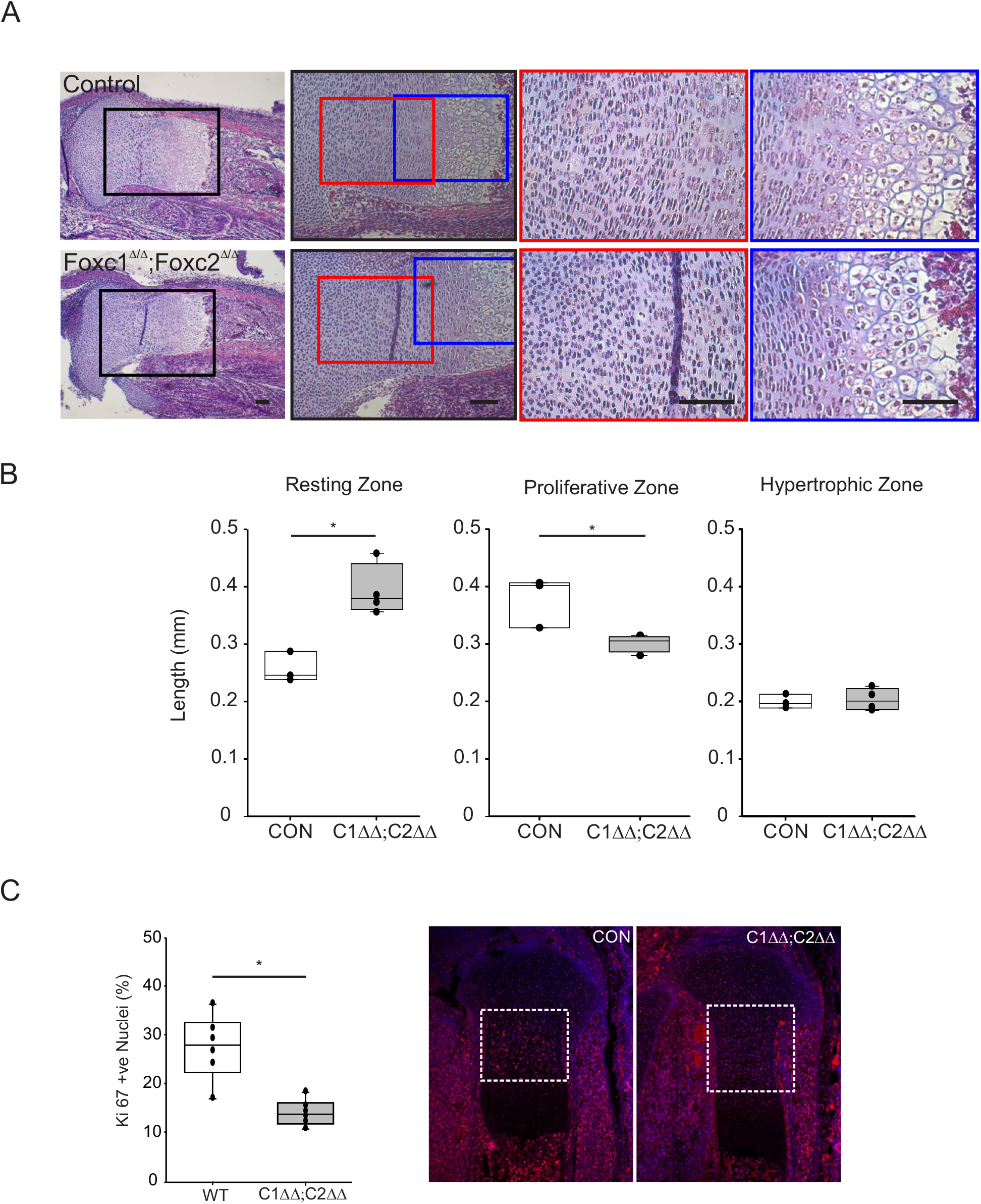
Disrupted growth plate organization in *Col2-cre;Foxc1^Δ/Δ^;Foxc2^Δ/Δ^* mutants. (A) Hematoxylin-Eosin staining of the 16.5 dpc proximal tibia growth plate. Enlarged sections are depicted with coloured boxes. (B) Length of the resting, proliferative and hypertrophic zone were measured in control (CON) or *Col2-cre;Foxc1^Δ/Δ^;Foxc2^Δ/Δ^* (C1*ΔΔ*;C2*ΔΔ*) growth plate. (C) Chondrocyte proliferation in control of *Col2-cre;Foxc1^Δ/Δ^;Foxc2^Δ/Δ^* was measured by Ki67 immunofluorescence.

### Endochondral ossification gene expression is reduced in Col2-cre;Foxc1^Δ/Δ^;Foxc2^Δ/Δ^ embryos

Given that *Foxc1* and *Foxc2* are transcription factors, it is expected that these genes exert their effects through regulation of gene expression. To further understand how *Foxc1* and *Foxc2* contribute to the formation of the skeleton we analysed gene expression of endocondral genes in *Col2-cre;Foxc1^Δ/Δ^;Foxc2^Δ/Δ^* embryos. We isolated RNA from the rib cage of 16.5 dpc embryos and monitored gene expression by quantitative reverse transcriptase-PCR (qRT-PCR). We chose the rib cage as a tissue source as non-skeletal tissues could be efficiently dissected away from the skeletal elements. Expression of *Foxc1* and *Foxc2* was reduced in rib RNA from *Col2-cre;Foxc1^Δ/Δ^;Foxc2^Δ/Δ^* embryos (Fig 8A). We assayed expression of genes that act throughout all stages of endochondral ossification and found that all genes assessed had decreased expression levels when *Foxc1* and *Foxc2* were absent. For example genes expressed early in the formation of chondrocytes (*Sox9, Sox6, and Col2a*) were reduced, while genes expressed in later chondrocyte differentiation stages such *Ihh, Fgfr3, ColX* as well as genes expressed during osteoblast formation and mineralization (*Sp7, Runx2, Col1a* and *Spp1*) were also reduced in *Col2-cre;Foxc1^Δ/Δ^;Foxc2^Δ/Δ^* rib RNA (Fig 8A).

**Figure 8.**
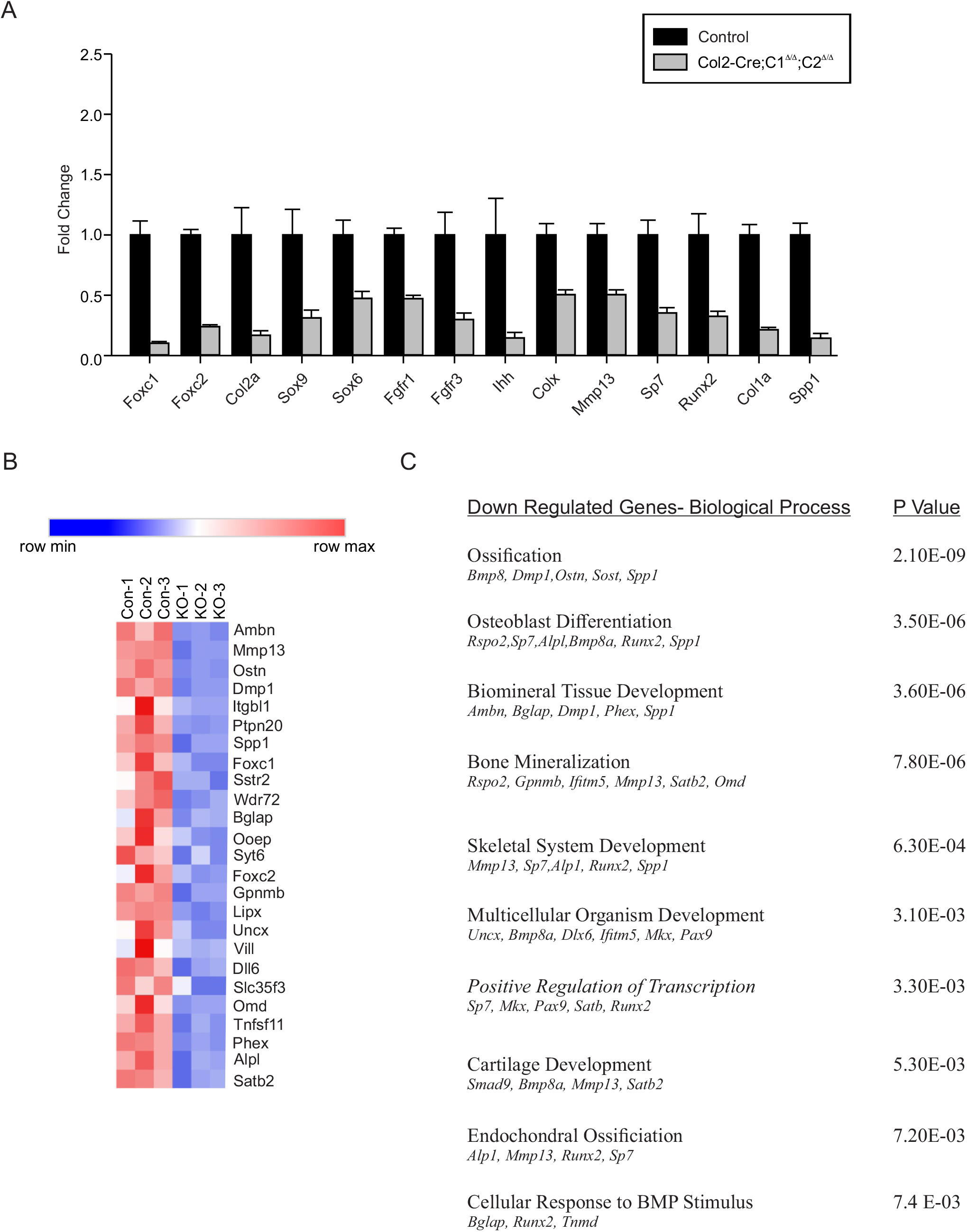
Loss of Foxc1 and Foxc2 function in chondrocytes results in a general reduction of mRNA levels of genes expressed throughout endochondral ossification. (A) RNA from the rib cage from control or *Col2-cre;Foxc1^Δ/Δ^;Foxc2^Δ/Δ^* embryos at 16.5 dpc was isolated and gene expression monitored by qRT-PCR. (B) Heat map of the top 25 genes down regulated in 16.5 dpc *Col2-cre;Foxc1^Δ/Δ^;Foxc2^Δ/Δ^* rib RNA as determined by RNA-seq analysis. (C) Functional annotation genes down regulated *Col2-cre;Foxc1^Δ/Δ^;Foxc2^Δ/Δ^* mutants.

To gain a complete picture of gene expression changes in response to deletion of *Foxc1* and *Foxc2* in chondrocyte progenitors, we performed RNA-seq from three additional samples of 16.5 rib cage RNA isolated from control and *Col2-cre;Foxc1^Δ/Δ^;Foxc2^Δ/Δ^* embryos. In total, we found 83 genes downregulated and 232 genes upregulated in rib cage RNA isolated from mutant embryos compared to controls (log2^foldchange^ +/− 1; p value <0.05; false discovery rate <0.01Supplemental Table 1 and 2). The 25 genes with the greatest reduction in expression in *Col2-cre;Foxc1^Δ/Δ^;Foxc2^Δ/Δ^* embryos (Fig 8B) included many of the genes involved in endochondral ossification whose expression. Gene ontology analysis revealed that the majority of biological functions affected in our down regulated gene set, were involved in endochondral ossification, including cartilage development, osteoblast differentiation and ossification (Fig 8C). Functional classification of the genes present in the upregulated data set were enriched with those involved in lipid metabolic processes, fatty acid metabolism and epidermis development (Supplemental Table 3). Together, data from these gene expression analyses suggest that loss of *Foxc1* and *Foxc2* function in *Col2-cre* expressing cells affected many stages of chondrocyte and osteoblast differentiation during the endochondral ossification processes.

Next we examined the expression of chondrocyte differentiation genes in formation of the growth plate of *Col2-cre;Foxc1^Δ/Δ^;Foxc2^Δ/Δ^* embryos in more detail. We found that the localization of SOX9 and SOX6 protein was unaffected in the tibia growth plate. These proteins were found throughout the resting zone and proliferative zone in the proximal tibia at 16.5 dpc (Fig 9A,B). Expression of *Fgfr3* was detected in the proliferative zone of *Col2-cre;Foxc1^Δ/Δ^;Foxc2^Δ/Δ^* embryos (Fig 9C) The extent of this expression domain appeared reduced in the *Col2-cre;Foxc1^Δ/Δ^;Foxc2^Δ/Δ^* embryos, likely owing to the reduced size of the proliferating zone (Fig 7). RUNX2 protein localized to the prehypertrophic chondrocytes of *Col2-cre;Foxc1^Δ/Δ^;Foxc2^Δ/Δ^* embryos (Fig 9D) and *Ihh* mRNA was also expressed in this region but expression intensity was reduced(Fig 9E). Localization of COLX protein in hypertrophic chondrocytes was also altered in the tibia growth plate of *Col2-cre;Foxc1^Δ/Δ^;Foxc2^Δ/Δ^* embryos at 16.5 dpc. Although the protein was present, its localization zone was expanded into the primary ossification in *Col2-cre;Foxc1^Δ/Δ^;Foxc2^Δ/Δ^* embryos (Fig 9F,H). MMP13 protein was also localized correctly to the hypertrophic chondrocytes at the osteochondral interface. Together these expression patterns suggest that in *Col2-cre;Foxc1^Δ/Δ^;Foxc2^Δ/Δ^* embryos. chondrocytes are able to correctly form and progress through their differentiation processes however this progression is altered as evidenced by expanded COLX protein.

**Figure 9.**
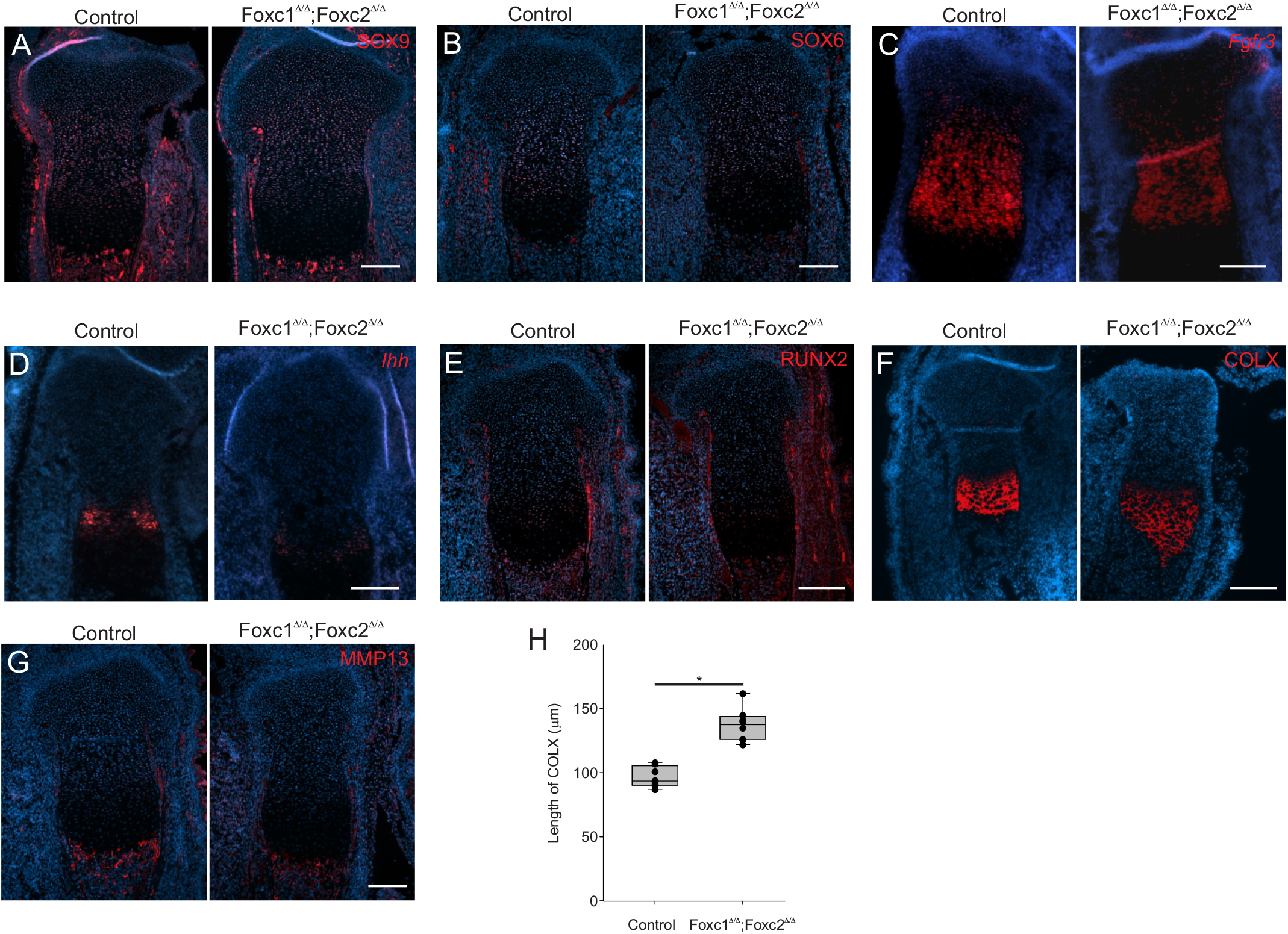
Altered chondrocyte differentiation in Col2-cre;Foxc1D/D;Foxc2D/D embryos. Gene expression patterns markers of chondrocyte differentiation were monitored in the proximal tibia growth plate at 16.5 dpc in control or *Col2-cre;Foxc1^Δ/Δ^;Foxc2^Δ/Δ^* embryos. (A) SOX9 immunofluorescence (IF) SOX6 IF (C) *Fgfr3* in situ hybridization (D) *Ihh* ISH (E) RUNX2 IF (F) COLX IF; (G) MMP13 IF. Length of RUNX2 (H) and COLX (I) positive cells in control vs *Col2-cre;Foxc1^Δ/Δ^;Foxc2^Δ/Δ^* growth plates. Scale bar 200 mm.

Finally we assessed the effect of loss of *Foxc1* and *Foxc2* function in the *Col2-cre* expressing cells on the ossification process. Von Kossa staining in the 16.5 dpc tibia indicated that ossification occured in the periosteum and osteochondral interface, however large areas of the bone marrow space were unmineralized (Fig 10 A, A’, B, B’). We next examined whether osteoblast formation occurred correctly in the *Col2-cre;Foxc1^Δ/Δ^;Foxc2^Δ/Δ^* embryos by monitoring Osterix (OSX) and Osteopontin (OPN) protein localization. In 16.5 dpc tibias, OSX and OPN positive cells were found in the osteochondral interface and in the primary ossification center (Fig 10C,D), indicating that osteoblasts formed in the *Col2-cre;Foxc1^Δ/Δ^;Foxc2^Δ/Δ^* embryos. Osteoblasts containing COL1a protein were detected at the osteochondral interface, the primary ossification center, the periosteum and groove of Ranvier in both control and *Col2-cre;Foxc1^Δ/Δ^;Foxc2^Δ/Δ^* embryos further indicated that osteoblast could form in the mutant bones (Fig 10 E,F). VEGFA protein localization was present but reduced in the hypertrophic chondrocytes in Col2-cre;Foxc1^Δ/Δ^;Foxc2^Δ/Δ^ embryos (Fig 10 G,H). Together these findings indicate that although endochondral bone formation and mineralization is abnormal in Col2-cre;Foxc1^Δ/Δ^;Foxc2^Δ/Δ^ embryos, the molecular processes that regulate this bone formation do occur in Col2-cre;Foxc1^Δ/Δ^;Foxc2^Δ/Δ^ embryos albeit in a disrupted manner.

**Figure 10.**
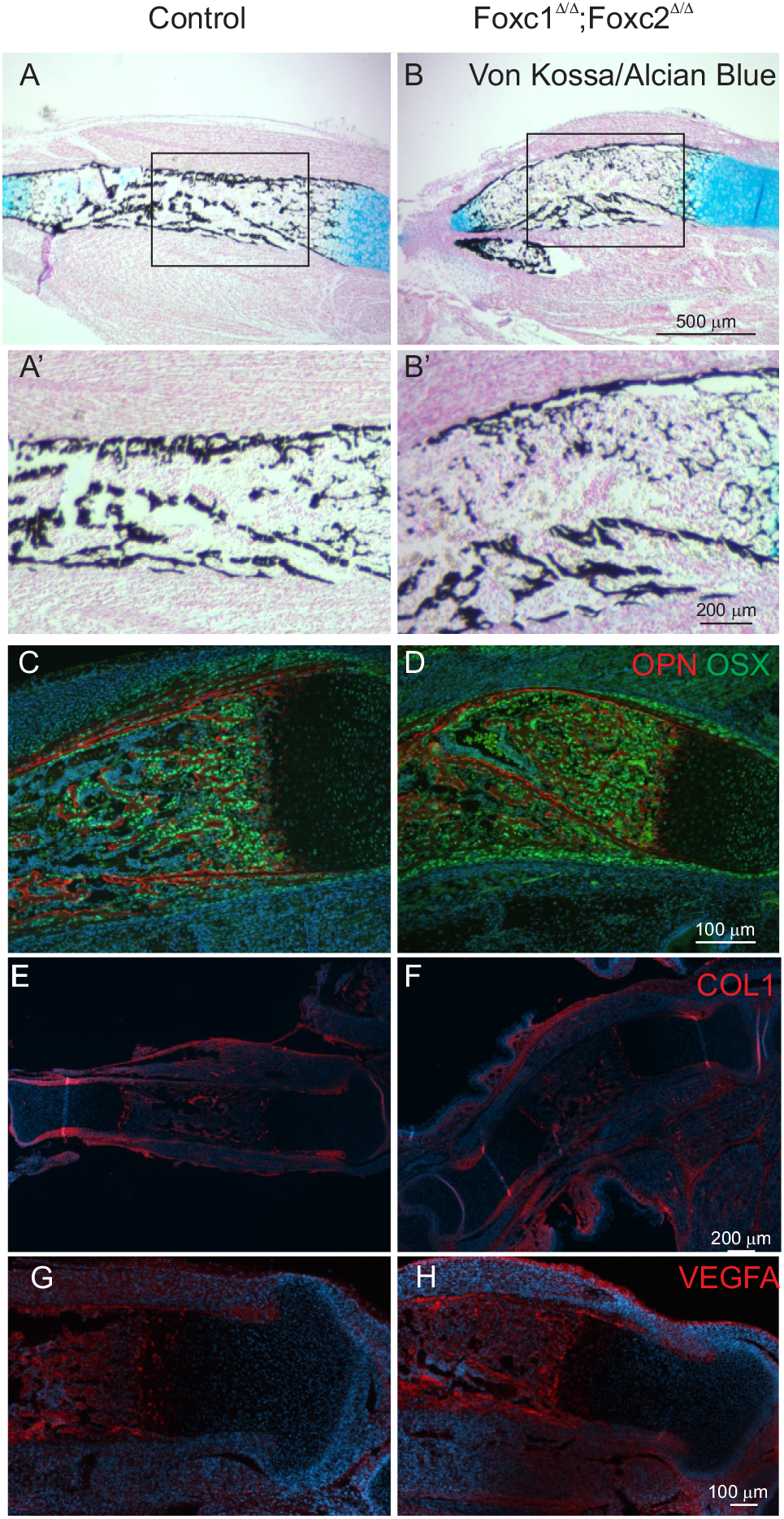
Impaired mineralization in *Col2-cre;Foxc1^Δ/Δ^;Foxc2^Δ/Δ^* mutants. (A,A’-B,B’) Mineralization in the primary ossification center of control and *Col2-cre;Foxc1^Δ/Δ^;Foxc2^Δ/Δ^* embryos determined by Von Kossa staining. (C-D) Levels of OSTEOPONTIN (Red) and OSTERIX protein (E,F,) COLLAGEN I and MMP13 localization (G-H) VEGFA localization.

## Discussion

*Foxc1* and *Foxc2* are required for normal skeletal development. We demonstrate that *Foxc1* and *Foxc2* are important regulators of chondrocyte formation and function required during endochondral ossification. *Foxc1* expression, but not *Foxc2*, is directly regulated by SOX9 activity. Enforced expression of *Foxc1* promoted the chondrocytic differentiation of mouse ES cells. Together these results indicate that *Foxc1* acts early in the formation of chondrogenic cells. This idea is further illustrated given that expression of *Foxc1* in the developing growth plate was enriched in the resting zone chondrocytes, with less mRNA detected in more differentiated cells. Loss of both *Foxc1* and *Foxc2* function in early Col2-expressing chondrocytes resulted in a disruption of endochondral ossification, leading to severe skeletal hypoplasia.

Little is known about the mechanisms that regulate *Foxc1* and *Foxc2* gene expression in the developing endochondral skeleton. We provide evidence that SOX9 can directly regulate expression of *Foxc1* in chondrocyte cells. Overexpression of *Sox9* in mES cells was sufficient to induce *Foxc1* mRNA. SOX9 binding to four distal regulatory regions in the Foxc1 gene had been suggested from ChIP-seq studies (29). We confirmed SOX9 association with regulatory elements in the *Foxc1* gene and ascribed functional activity to these regions. Although SOX9 was able to bind to all four elements, only one element (Distal C) elicited a transcription response to SOX9 in ATDC5 chondrocyte cells. A second region (Distal B) did confer increased reporter activity in ATDC5, suggesting it contains a regulatory information to activate Foxc1 expression in chondrocyte cells, however its activity is likely independent of SOX9 as its activity was unaffected by SOX9 levels. Recently, expression of *FOXC1* mRNA in human breast cancer cells was controlled by SOX9 (34). Regulation of *Foxc1* by SOX9 is further supported by the reduction of *Foxc1* expression levels in the limbs of *Sox9* deficient embryos (17). This study also reported a decrease in *Foxc2* mRNA in *Sox9* mutant mice, although we did not detect any induction of *Foxc2* expression using Sox9-inducible mES cells. It should be noted that these cells were not differentiated towards the chondrocyte lineage and therefore it is possible that SOX9 might regulate *Foxc2* expression in a tissue-specific context. *Foxc1* activation by SOX9 suggests a role for *Foxc1* function in the early stages of chondrocyte differentiation. We observed that *Foxc1* expression is spatially enriched in resting zone chondrocytes compared to proliferating and hypertrophic zone chondrocytes which represent later differentiation stages. As well, over-expression of *Foxc1* in mouse ES cells led to an increased chondrocyte differentiation capacity, suggesting this early increase in Foxc1 levels promoted cells to the chondrocyte fate.

We also demonstrate the comparative expression patterns of *Foxc1* and *Foxc2* in the developing mouse limb. We determined that *Foxc1* and *Foxc2* have some overlapping as well as distinct expression patterns in the growing limb. Expression of *Foxc1* and *Foxc2* mRNAs were predominantly detected in the perichondrium surrounding the growth plate and in the resting zone chondrocytes. Although much of the overall expression pattern in these regions for *Foxc1* and *Foxc2* was similar, there were areas where only *Foxc1* or *Foxc2* expression could be detected, suggesting these factors may not have completely overlapping functions. Very few *Foxc1* or *Foxc2* expressing cells could be observed in the proliferating zone, pre- and hypertrophic chondrocytes at 16.5 dpc. This expression pattern suggests that the chondrodysplasia that we observed in the *Col2-cre;Foxc1^Δ/Δ^;Foxc2^Δ/Δ^* mutants arose from either an indirect effect of *Foxc1* and *Foxc2* acting in the perichondrium to regulate chondrogenesis or as a consequence of loss of *Foxc1* and *Foxc2* function at an early stage of chondrocyte differentiation that disrupts cell function throughout later stages of development. We did detect *Foxc1* expression in a subset of cells lying between the perichondrium and the proliferating chondrocytes, suggesting that *Foxc1* may be expressed in borderline chondrocytes that supply skeletal stem cells later in life (35). Spatially, we observed increased Foxc1 and Foxc2 hybridization signals in distal skeletal elements compared to proximal ones. This expression pattern may reflect a role for Foxc1 and Foxc2 in the proximal-distal patterning of the limb. Additionally, as endochondral ossification in the limb proceeds in a proximal to distal manner, this expression pattern may reflect a role for Foxc1 and Foxc2 in the early stages in the formation of skeletal elements and expression decreases as the element matures.

Global knock out of either *Foxc1* or *Foxc2* results in a number of skeletal phenotypes in mice (19, 20, 22). These mutations predominantly affect the axial skeleton, although the bones of the appendicular skeleton do display a modest reduction in size. Loss of both *Foxc1* and *Foxc2* in *Col2-cre* expressing cells, results in a more severe disruption to the endochondral skeleton compared to loss of either *Foxc1* or *Foxc2* alone. We observe a general disorganization of the growth plate that affects endochondral ossification processes as indicated by the broad reduction in gene expression of regulators of these events. We reported a reduction in gene expression affecting all stages of chondrocyte differentiation and function as well as genes involved in osteoblast formation and mineralization. The growth plate of the tibia was disorganized in the *Col2-cre;Foxc1^Δ/Δ^;Foxc2^Δ/Δ^* mutants, and the columnar arrangement of the proliferative chondrocytes did not form, resulting in decreased cell proliferation that likely accounts for the reduced bone size. We did observe a reduction in *Ihh* mRNA levels which likely accounts for the impaired proliferation observed in *Col2-cre;Foxc1^Δ/Δ^;Foxc2^Δ/Δ^* mutants. Chondroprogenitor cells lacking *Sox9* display a reduction in the length of the columnar chondrocyte zone and a reduced cell proliferation (15). Moreover, Sox6 function is required to maintain columnar organization of proliferative zone chondrocytes (12). We observe similar phenotypes in *Col2-cre;Foxc1^Δ/Δ^;Foxc2^Δ/Δ^* embryos although SOX9 and SOX6 protein levels were unaffected, suggesting that *Foxc1* and *Foxc2* may function in common aspects of *Sox9*/*Sox6*-dependent chondrogenic processes.

We did observe some differences in the *Col2-cre;Foxc1^Δ/Δ^;Foxc2^Δ/Δ^* mutants compared with global *Foxc1* or *Foxc2* gene mutations. Germline deletion of *Foxc2* results in fused ribs (20). The number and positioning of the ribs was not affected in our mutants, suggesting a role for *Foxc2* in rib patterning and specification that occurs prior to rib chondrocyte formation. The sternum does not fully minerize in *Foxc1*^−/−^ mice and completely lacks ossification of the xiphoid process (19). In the *Col2-cre;Foxc1^Δ/Δ^;Foxc2^Δ/Δ^* mutants the ribcage is reduced in size but mineralization along the sternum appears unaffected. This difference in phenotype may arise either from a less efficient Cre-activity in cells that form the sternum or that a portion of these cells that make up this structure arise from cells that do not express *Col2a*. COLX and MMP13 protein levels were also markedly reduced in germline *Foxc1^−/−^* mutants (21) whereas COLX expression was expanded in the *Col2-cre;Foxc1^Δ/Δ^;Foxc2^Δ/Δ^* mutants and MMP13 levels were not affected. These observed phenotype differences may reflect a function for *Foxc1* and *Foxc2* in progenitor cells prior to the onset of *Col2a* expression or in cells that do not express *Col2a* and thus did not delete Foxc1 and Foxc2.

The combined loss of both *Foxc1* and *Foxc2* in Col2a-expressing cells resulted in delayed endochondral ossification events in both the appendicular and axial skeleton. These disruptions were more pronounced in the axial skeleton. There are a number of reasons to account for such observations. First, Col2-cre is detected in the sclerotome of the somites prior to these cells migrating to and condensing at sites of future vertebral bone formation (36). In contrast Col2-cre activity is not detected in the limbs until mesenchyme cells have migrated and condensed where the future bones will develop. This suggests that Foxc1 and Foxc2 are deleted at an earlier stage of vertebral bone formation that may prevent formation of adequate numbers of sclerotome cells needed to form the axial skeleton. Second, this disruption may reflect a difference in the chondrogenesis process occurring in the axial skeleton compared to the appendicular skeleton. In the sclerotome, cells must first receive Sonic Hedgehog signal from the notochord in order to initiate chondrogenesis, whereas such a signalling is not required in the limb skeleton (37, 38). It is possible that *Foxc1* and *Foxc2* may function in processing this SHH signal needed for chondrogenesis in the sclerotome. Third, additional transcription factors present in the limb skeleton may compensate for loss of *Foxc1* and *Foxc2* and such compensation does not occur in the axial skeleton. Experiments to address such mechanisms are being pursued in our laboratory.

In conclusion, *Foxc1* and *Foxc2* functions in chondrocytes are required for correct endochondral ossification events to occur. Loss of *Foxc1* and *Foxc2* function in Col2-cre expressing cells results in skeletal dysplasia and disrupts skeletal mineralization. The expression patterns for these factors in the growth plate suggest they act at early stages of chondrogenesis and their loss of function impacts later chondrogenic differentiation stages. Chondrocytes do form in the absence of *Foxc1* and *Foxc2* but they are unable to correctly differentiate resulting in disorganized growth plate, reduced chondrocyte proliferation and delays in chondrocyte hypertrophy. Such disruptions have the overall effect of preventing correct ossification of the endochondral skeleton.

## Experimental Procedures

### Cell culture and in vitro differentiation

Mouse embryonic stem (mES) cells containing an inducible Sox9 or Foxc1 expression cassette (27, 30) were obtained from either the Coriell Institute (Sox9) or Dr. Minoru Ko (Foxc1). Mouse ES cells were cultured in DMEM media (MilliporeSigma) containing 15% Fetal Bovine Serum (FBS; Gibco), 1% L-Glutamine (Life Technologies), 1 mM β-mercaptoethanol (MilliporeSigma), 0.1 mM Non-Essential Amino Acids (Lifetechnologies), 1000 units/ml Leukemia Inhibitory Factor (MilliporeSigma), 1% Penicillin/Streptomycin (Lifetechnologies), and 1μg/ml Doxycycline (DOX). To induce gene expression, cells were repeatedly washed with PBS before replacement with DOX-free media.

ATDC5 cells were purchased from European Collection of Authenticated Cell Cultures and cultured in DMEM:F12 containing 5% fetal bovine serum. Mutagenesis of the Foxc1 gene was achieved by using Alt-R CRISPR-Cas9 system (Integrated DNA Technologies) using a predesigned gRNA (5’-CAACATCATGACGTCGCTGC-3’) that targeted the second helix of the FOXC1 Forkhead box DNA- binding domain. Ribonucleoprotein complexes were transfected into ATDC5 cells using RNAiMAX. Two days post-transfection, cells were then plated at a single cells per well of 96 well plate by diluting cells to approximately 80 cells/ml and 100 μl cell solution was added to each well. Single cells were then expanded to larger culture volumes and were screened for a reduction in Foxc1 protein levels by immunoblotting with anti Foxc1 antibodies (Origene), followed by sequencing of the Foxc1 ORF. Two cell lines were selected (crFOXC1-1 and crFOXC1-8) and used for subsequent functional studies. Both lines displayed similar properties in chondrocyte differentiation experiments but results from a single line (crFOXC1-1) were reported here.

### RNA-isolation and qRT-PCR

RNA was isolated from cell cultures and tissues using RNeasy mini kit (Qiagen) following the manufacturer’s protocol. All lysates were homogenized using QIAshredder (Qiagen). Tissue lysates were first disrupted with a microcentrifuge pestle before passing through a QIAshredder. RNA (500 ng) was then reverse transcribed to cDNAs using the QuantiTect reverse transcription kit (Qiagen). qRT-PCR analysis was performed as described previously (39) (40). All qRT-PCR experiments were performed with at least three biological replicates and each contained three technical replicates. Primers used for analysis were obtained as pre-designed PrimeTime qPCR Primer Assays (Integrated DNA Technologies, IDT).

### Chromatin Immunoprecipitation (ChIP)assays

ChIP assays were performed as described previously (41) with the following modifications. Chromatin from ATDC5 cells sheared in ice using Bronson Sonifier (10 cycles at 30% amplitude for 30 s with a 60 s rest).

Cross-linked chromatin extracts were incubated overnight with 2 μg anti-SOX9 antibody (Millipore), Acetylated Histone H3 (Millipore), or rabbit IgG. Amplification of recovered chromatin was performed by PCR using the following primers. Col2a intron 1 forward primer (5’-TGA AAC CCT GCC CGT ATT TAT T -3’) reverse primer (5’-GCC TTG CCT CTC ATG AAT GG-3’). Foxc1 Distal A forward forward primer (5’-GCC CTG AAT CCA GAA ACT TG -3’) reverse primer (5’-GCG AAT TCA TAT GGT TTT TCC -3’). Foxc1 Distal B forward primer (5’-GGCCATCATGTCTAGGGGAA -3’) reverse primer (5’-GTTGCTCTGAACTTGGGGTG -3’). Foxc1 Distal C forward primer (5’- TGT GAA ATC GCC TGT GAG AG-3’) reverse primer (5’- -CCC CAT ATC CTC TTT GAG AGC3’). Foxc1 Distal D forward primer (5’TGT CAG GAG AAC TGC TGT AAG AA- -3’) reverse primer (5’-CTC TAG GCT GAC CAC GCT GT -3’).

### Reporter cloning and luciferase assays

DNA fragments corresponding to mouse Foxc1 regulatory regions Distal A (mm10 chr13:31,764,541-31,764,717), Distal B (mm10 chr13:31,765,465-31,765,623), Distal C (mm10 chr13:31,779,560-31,779,803) and Distal D (mm10 chr13:31,820,626-31,820,791) synthesized as gBlock fragments (IDT) and were cloned into the Eco RV site of pGL4.23-luc2/minP vector (Promega) using Gibson Assembly. Plasmids containing the correct regulatory sequence were confirmed by sequencing. Dual luciferase reporter assays (Promega) were performed as described previously (40)

### Chondrocyte differentiation procedures

For chondrocyte differentiation experiments, mES cells were grown in DOX-free media for 2 days prior to induction of a chondrocyte differentiation protocol as described in ref ((31)). Chondrocyte differentiation of ATDC5 cells was initiated by supplementing cell culture media with 1x Insulin, Transferrin and Selenium supplement (Cellgro). Differentiation media was replenished every 2-3 days.

### Mouse models

All research using mouse models was approved by the University of Alberta Animal Care and Use Committee (AUP804). *Col2-cre* mice (32) were kindly provided by Dr René St-Arnaud (Shriners Hospital for Children, Montreal). Foxc1^fl/fl^;Foxc2^fl/fl^ (33) were crossed with *Col2-cre^+/−^* mice. Timed pregnancies were conducted by crossing male *Col2-cre^+/−^;Foxc1^+/fl^;Foxc2^+/fl^* mice to female *Foxc1^fl/fl^;Foxc2^fl/fl^* mice. Mice were maintained on mixed background C57B6 (Col2-cre) and Black Swiss (*Foxc1^fl/fl^;Foxc2^fl/fl^*) Crosses were set up in the afternoons and vaginal plugs were monitored in the morning, with noon of the day a positive plug was detected designated as 0.5 dpc. All comparisons were made between littermates. Weaned mice were genotyped using ear notch biopsies, while embryos were genotyped using skin DNA. Genotyping was performed using the KAPA mouse genotyping kit (MilliporeSigma). The following primer pairs were used: Foxc1 (forward 5’-ATTTTTTTTCCCCCTACAGCG-3’; reverse 5’-ATCTGTTA GTATCTCCGGGTA-3’), Foxc2 (forward 5’-CTCCTTTGCGTTTCCAGTGA -3’; reverse 5’-ATTGGTCCTTCGTCT TCGCT -3’) and Col2-cre (forward 5’-GCCTGCATTACCGGTCGATGCAACGA -3’; reverse 5’-GTGGCAGATGGCGCGGCAACACCATT -3’).

### Skeletal preps and analysis

Skeletons from embryos collected at 18.5 dpc were stained with Alizarin Red and Alcian Blue as described in (42). Images were captured with an Olympus E520 digital camera.

### In situ hybridization

Embryos were collected at the desired stage and fixed in 4% paraformaldehyde overnight at 4 °C. Tissues were washed in PBS overnight prior to embedding in paraffin. Sections (7 um) were collected onto superfrost slides (Fisher Scientific). In situ hybridization was performed using the RNAscope multiplex in situ hybridization kit (Advanced Cell Diagnostics). The following probes were used. Foxc1 (412851-C2, lot 19155B); Foxc2 (406011, lot 18289A); Fgfr3 (440771, lot 18289A); and negative control probe (310043; lot 18197A).

### Immunofluorescence

The following antibodies were used for immunofluorescence microscopy: COL IIa (Abcam, ab185430, lot GR3320839, 1:100); COLX (Abcam, ab5832, lot GR3210868-2, 1:50); COL I (Abcam, ab88147; GR3225500-1, 1:100); MMP13 (Abcam, ab39012, lot GR157414-15 1:100); RUNX2 (Abcam, ab76956, lot, 1:200) SOX6 (Abcam, ab30455, lot GR3174880-1, 1:1000); VEGFA (Abcam, ab1316, lot GR3200812. 1:100), SOX9 (MilliporeSigma, AB55535, lot 3282152, 1:200); OPN (SCBT, sc22536-R, lot C2307, 1:100); OSX (SCBT, sc21742, lot D2908, 1:100); Ki67 (Bethyl, IHC00075, 1:100). Antibodies against GFP were a generous gift from Dr Luc Berthiume (University of Alberta) and were used at a dilution of 1:500.

Paraffin sections were collected as described for in situ hybridization procedures. Slides were baked at 70 °C for 1 hour followed by rehydration through xylene and graded ethanol series. For SOX9, SOX6, OSX, OPN, KI67, RUNX2 and GFP antibodies antigen retrieval was conducted boiling slides in citrate buffer (10 mM trisodium citrate, pH 6.0; 0.05% Tween20) for 20 minutes. For COL1, COL2a, COLX, MMP13, VEGFA antibodies, sections were incubated in hyaluronidase. After antigen retrieval slides were blocked in 5% donkey serum in PBS with 0.05% Triton X-100 (PBSX) for one hour followed by incubation with primary antibody overnight at 4°C. Slides were washed in PBSX before incubation in secondary antibody for 1 hour at room temperature, followed by staining with DAPI and mounted with coverslips using Prolong Gold.

### RNA-seq

RNA was isolated from the ribs of three control and three *Col2-cre;Foxc1^Δ/Δ^;Foxc2^Δ/Δ^* 16.5 dpc embryos as described above. RNA Sample quality control was performed using the Agilent 2100 Bioanalyzer. Samples with RNA Integrity Number > 8 were used for library preparation following the standard protocol for the NEBnext Ultra ii Stranded mRNA (New England Biolabs). Library construction and sequences was carried out at the Biomedical Research Centre Sequencing Facility (University of British Columbia, Canada). Sequencing was performed on the Illumina NextSeq 500 with Paired End 42bp × 42bp reads. De-multiplexed read sequences were then aligned to the reference sequence using STAR (https://www.ncbi.nlm.nih.gov/pubmed/23104886) aligners. Two different pipelines, HISAT2-featureCounts-DESeq2 and STAR-RSEM-DESeq2 were used for the downstream analysis through an inhouse script. For both the pipelines, mouse reference genome GRCm38 was downloaded from NCBI (https://www.ncbi.nlm.nih.gov/assembly/GCF_000001635.20/) while gene annotation was downloaded from gencode (https://www.gencodegenes.org/mouse/). Briefly, read alignment was performed using both HISAT2 (https://pubmed.ncbi.nlm.nih.gov/25751142/) and STAR (https://www.ncbi.nlm.nih.gov/pubmed/23104886). Quantification using featureCounts (https://pubmed.ncbi.nlm.nih.gov/24227677/) and RSEM (https://pubmed.ncbi.nlm.nih.gov/21816040/) while Differential Expression Analysis was performed using DESeq2 (https://pubmed.ncbi.nlm.nih.gov/25516281/) for both. The final output was a list of genes with significant p-value (<0.05) after correction for multiple testing with a false discovery rate of less than 0.1. A combined list was obtained by averaging the genes with a filtered log2 fold change ± 1. Only common genes with consistent differential expression log2 fold change values between both the pipelines were included in the list while the inconsistent ones were excluded. Correlation for each sample between the pipelines was calculated using linear regression.

