## Supplemental Table 3 for "Loss of Foxc1 and Foxc2 function in chondroprogenitor cells disrupts endochondral ossification"

| Upregulated Genes Biological Processes | P-value |
|---|---|
| Peptide Cross-linking<br>*Crct1, Lce1a1, Lce1a2, Lce1b,Sprr2d* | 3.1E-18 |
| Keratinocyte Differentiation<br>*Crct1, Lce1a1, Lce1a2, Lce1b,Sprr2d* | 2.8E-15 |
| Lipid Metabolic Process<br>*Agpat5, Hmgcs2, Angptl8, Apod, Cpt1b, Cidea, Cyp11a1, Ephx2, Far2, Pnpla2* | 9.0E-8 |
| Epidermis Development<br>*Flg. Lce1f, Lce1h, Ptch2, Sprr2d* | 2.8E-8 |
| Oxidation-Reduction Process<br>*Ddo, Chdh, Cyp11a1, Far2, Fmo2, Sod3, Tryp1* | 7.9E-4 |
| Triglyceride Catabolic Process<br>*Lipe, Pnpla2, Pnpla3, Plin1* | 8.7E-4 |
| Regulation of Lipid Metabolic Process<br>*Angptl8, Hnf4a, Irs4, Pparg,* | 3.8E-3 |
| Brown Fat Cell Differentiation<br>*Adipoq, Fabp4, Mrap, Pparg* | 4.1E-3 |
| Negative Regulation of Smooth Muscle Cell Proliferation<br>*Ndrg2, Adipoq, Apod, Pparg* | 5.7E-3 |
| Positive Regulation of Fatty Acid Biosynthetic Process<br>*Mixipl, Agt, Hnf4a* | 7.7E-3 |
